# Neural correlate of reduced respiratory chemosensitivity during chronic epilepsy

**DOI:** 10.1101/2022.01.06.475212

**Authors:** Amol Bhandare, Nicholas Dale

## Abstract

Central autonomic cardiorespiratory dysfunction underlies sudden unexpected death in epilepsy (SUDEP). Here we used single cell neuronal Ca^2+^ imaging and intrahippocampal kainic acid (KA)-induced chronic epilepsy in mice to investigate progressive changes in key cardiorespiratory brainstem circuits during chronic epilepsy. Following induction of status epilepticus (SE), adaptive ventilatory responses to hypercapnia were reduced in mice with chronic epilepsy for 5 weeks post-SE with partial recovery at week 7. These changes were paralleled by post-SE alterations in the chemosensory responses of neurons in the retrotrapezoid nucleus (RTN). Neurons that displayed adapting responses to hypercapnia were less prevalent and exhibited smaller responses over weeks 3-5, whereas neurons that displayed graded responses to hypercapnia became more prevalent by week 7. Over the same period, chemosensory responses of the presympathetic rostral ventrolateral medullary neurons showed no change. Mice with chronic epilepsy showed enhanced sensitivity to seizures, which can invade the RTN and possibly impair further the chemosensory circuits. Our work suggests that assessment of respiratory chemosensitivity may have potential for identifying people at risk of SUDEP.

## Introduction

Every year 0.4-2/1,000 of people with epilepsy and 4-9/1,000 of people with drug-resistant epilepsy die as a result of sudden unexpected death in epilepsy (SUDEP) (Tomson *et al*., 2008; Ryvlin *et al*., 2013). There is currently no treatment or biomarker test available to identify people that are at higher risk of SUDEP. Although studies show that SUDEP mainly occurs in people with drug-resistant and generalised tonic-clonic seizures (Tomson *et al*., 2008; Ryvlin *et al*., 2013), it is still unclear why these seizures are more fatal in people with a history of seizures (Shorvon & Tomson, 2011). It is still unknown whether the incidence of SUDEP is higher in people with frequent seizures because each seizure has a fixed and finite chance of precipitating death or because there are cumulative changes in central cardiorespiratory circuitry resulting from history of seizures that progressively increase the risk of death. There is some evidence that cumulative autonomic cardiorespiratory changes due to repeated seizures might be a potential cause of SUDEP (Patodia *et al*., 2022).

Central autonomic cardiorespiratory brainstem circuits generate the rhythms for breathing and the sympathetic and parasympathetic activity that controls the heart (Spyer & Gourine, 2009; Guyenet, 2014). Evidence from epilepsy monitoring units (So *et al*., 2000; Tomson *et al*., 2008; Ryvlin *et al*., 2013; Sivathamboo *et al*., 2020) and *in vivo* rodent findings (Aiba & Noebels, 2015; Jefferys *et al*., 2019) implies that apnoea (either centrally mediated or obstructive) and bradycardia occurring immediately after seizures leads to SUDEP. Numerous studies have proposed different mechanisms of SUDEP including parasympathetic hyperactivity (Kalume *et al*., 2013), deficiency of Kv1.1 potassium channels (Glasscock *et al*., 2010; Moore *et al*., 2014; Aiba & Noebels, 2015), reduced heart rate variability (Surges *et al*., 2009), cardiac arrhythmia (Naggar *et al*., 2014; Bhandare *et al*., 2015; Stewart, 2018), hypoxemia (Bateman *et al*., 2010), hypercapnia (Seyal *et al*., 2010), pulmonary oedema (So *et al*., 2000), airway obstruction (Stewart *et al*., 2017; Jefferys *et al*., 2019; Irizarry *et al*., 2020), amygdala seizures (Park *et al*., 2020) along with role of neurotransmitter serotonin (Faingold *et al*., 2011; Buchanan *et al*., 2014), adenosine (Faingold *et al*., 2016; Shen *et al*., 2022) and loss of brainstem volume (Mueller *et al*., 2014) or glial cells (Patodia *et al*., 2019). The origins of SUDEP are likely to lie in the dysfunction of key brainstem circuits (So *et al*., 2000; Ryvlin *et al*., 2013; Mueller *et al*., 2014). In genetic mouse models of epilepsy, seizure-induced spreading depolarisation (recorded in the dorsal medulla) silences the brainstem respiratory neuronal network, and this has been proposed to cause SUDEP (Aiba & Noebels, 2015). While this suggests disruption or silencing of key cardiorespiratory outputs from the brainstem, it does not give insight into the specific neuronal circuitry and mechanisms involved.

A recent study has shown reduced ventilatory responses to hypercapnic challenges up to 30 days after pilocarpine-induced status epilepsy (SE) in rats (Maia *et al*., 2020). Additionally, lower hypercapnic ventilatory responses in people with epilepsy have been correlated with higher postictal transcutaneous CO_2_ rise following generalized convulsive seizures (Sainju *et al*., 2019). As the drive to breathe is critically dependent on PCO_2_, reductions in sensitivity to PCO_2_/pH of central chemosensory neurons might plausibly contribute to central apnoea and ultimately SUDEP. These chemosensory neurons are located at the ventral surface of the medulla and regulate breathing to maintain systemic PCO_2_ within physiological limits (Spyer & Gourine, 2009; Guyenet, 2014). Cardiovascular function is controlled by excitatory rostral ventrolateral medullary (RVLM) neurons in the brainstem (Spyer & Gourine, 2009; Guyenet, 2014). To address whether progressive changes in epilepsy alter the neuronal firing patterns and chemosensitivity in these critical brainstem networks, we used intravital microscopy to record from these brainstem nuclei in freely behaving mice (Bhandare *et al*., 2022).

By using genetically-encoded Ca^2+^ indicators and gradient refractive index (GRIN) optic fibres, to allow the direct visualization of dynamic cellular activity of deep brain cardiorespiratory neurons in awake mice at single cell resolution, we have recorded how seizures alter the activity and responses of chemosensitive respiratory RTN and cardiovascular RVLM neurons. We found that mice with chronic epilepsy (a state characterized by unprovoked recurrent seizures), compared to their pre-epileptic state, have reduced baseline respiratory activity (tidal volume, V_T_) for 5 weeks after induction of epilepsy. In addition, the adaptive breathing changes to hypercapnia were substantially weakened for 5 weeks following SE and showed partial recovery at week 7. By simultaneously recording neuronal activity from the same mice before and following the establishment of SE for several weeks, we have documented parallel changes in the chemosensory responses of RTN neurons that could explain the observed changes in the chemosensory control of breathing. Although this does not demonstrate a mechanism of SUDEP, we propose that the reduced chemosensitivity of RTN neurons, by lessening the baseline drive to breathe and reducing adaptive ventilatory responses to hypercapnia, could be a contributing mechanism of SUDEP.

## Results

### Inclusion/exclusion criteria for cells in the study

Following assessment for inclusion/exclusion criteria for imaging of both RTN and RVLM mice (Figure 1A), data from 15 mice comprising recordings from 146 (pre-SE), 96 (week-3 post-SE), 60 (week-5 post-SE) and 82 (week-7 post-SE) cells were included for analysis (Figure 1B). As the mice were unrestrained and able to move freely during the recordings, to visualize physical movements during Ca^2+^ analysis, high-definition videos of mice were synchronized to the simultaneous plethysmograph and Inscopix Ca^2+^ imaging recordings in Inscopix Data Processing Software and Spike2. Our recordings were performed at a single wavelength (Figure 1A). Therefore, it was essential to verify that any signals result from changes in Ca^2+^ rather than from movement artefacts.

**Figure 1:**
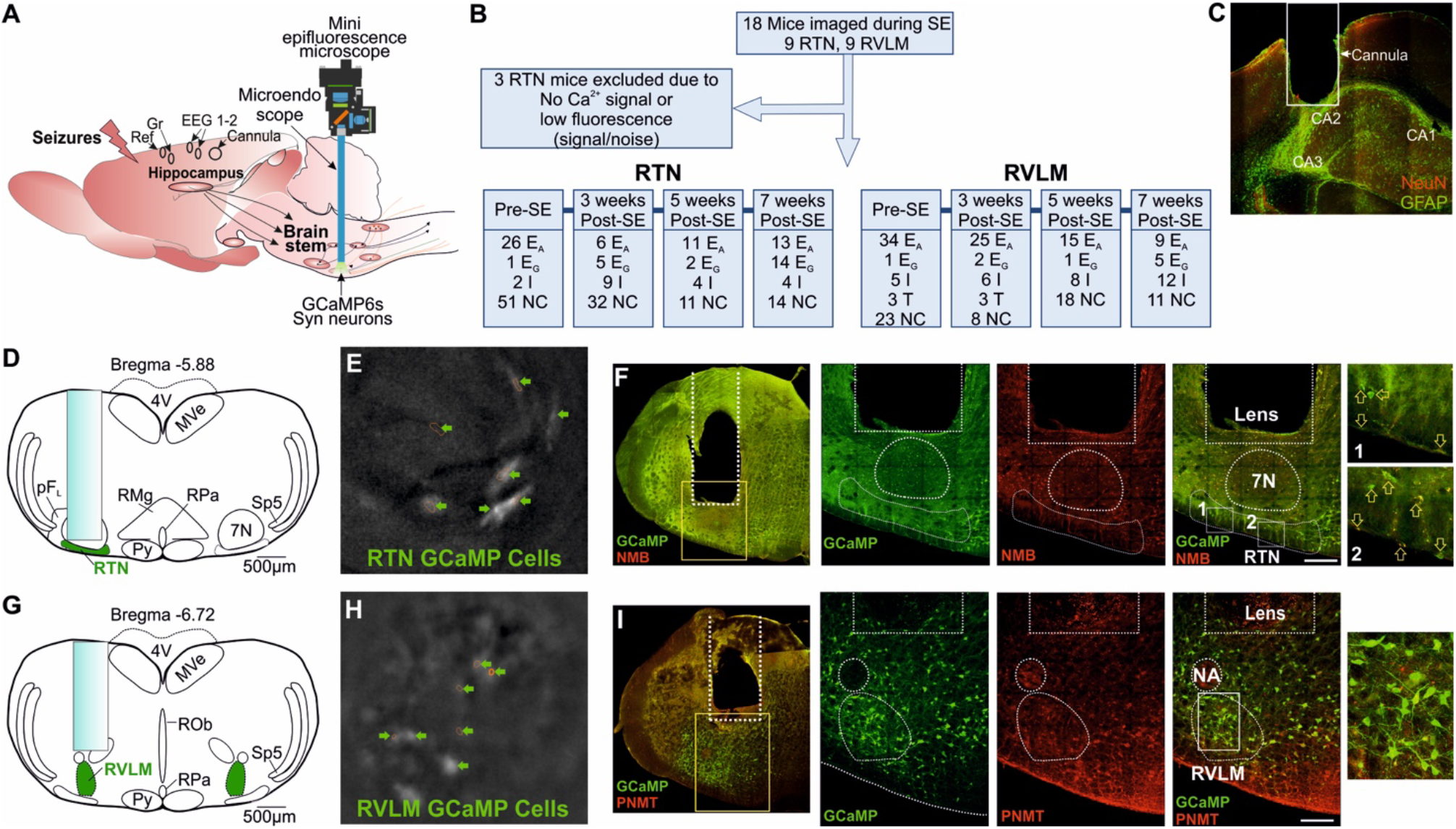
Microendoscopic experimental approach to target specific brainstem nuclei. A) Representation of GRIN lens (microendoscope), baseplate, and mini epifluorescence camera placement for recording of brainstem nuclei with hippocampal cannula for seizure induction and EEG (recording, reference and ground) electrodes. B) A CONSORT flow diagram for inclusion/exclusion of experiments and cells classification. Definitions of the range of firing patterns in response to hypercapnia are described in the text. C) Micrograph of hippocampal cannula placement for injection of KA. D) AAV9-Syn-GCaMP6s injection into the RTN with the lens position. E) GCaMP6s fluorescence of transduced RTN neurons in freely behaving mice. F) Micrograph of lens placement and viral transduction of neurons (green) relative to the facial nucleus (7N) and NMB+ RTN neurons (red). G) AAV9-Syn-GCaMP6s injection into the RVLM with the lens position. H) GCaMP6s fluorescence of transduced RVLM neurons in freely behaving mice. I) Micrograph of lens placement and viral transduction of neurons (green) and PNMT+ RVLM neurons (red). Scalebar in panel F and I represent 200µm; Abbreviations: E_A_, adapting; NC, non-coding; I, inhibited; E_G_, graded; T, tonic: 7N, facial motor nucleus; Py, pyramidal tract; MVe, medial vestibular nucleus; sp5, spinal trigeminal nucleus: RTN, retrotrapezoid nucleus; RMg, raphe magnus; RPa, raphe pallidus; ROb, raphe obscurus; pFL, parafacial lateral region; RVLM, rostral ventrolateral medulla; NA, nucleus ambiguus; NeuN, neuronal nuclei; GFAP, glial fibrillary acidic protein; NMB, neuromedin B; PNMT, phenylethanolamine N-methyltransferase.

Detailed procedures to assess and compensate for movement artefact have been given previously (Bhandare *et al*., 2022). In brief, if there was too much uncompensated motion artefact, so that we could not clearly analyze Ca^2+^ signals, we excluded these recordings from quantitative analysis. GCaMP6 signals were only accepted for categorization if the following criteria were met: 1) the features of the cell (e.g. soma, large processes) could be clearly seen; 2) they occurred in the absence of movement of the mouse or were unaffected by mouse movement; 3) fluorescence changed relative to the background; 4) the focal plane had remained constant as shown by other nonfluorescent landmarks (e.g. blood vessels). To assess the role of movement, we examined the GCaMP6 fluorescence in conjunction with mouse’s body movement. We included the cells that that did not evoke noticeable change in GCaMP6 fluorescence with the body movements and characterized by a fast rise and exponential decay in the absence of any movement as well as during movement.

### Seizures spread into the central autonomic cardiorespiratory network

Using methods that we have recently developed (Bhandare *et al*., 2022), neurons of the RTN and RVLM were transduced with an AAV to drive expression of Ca^2+^ reporter GCaMP6 under the control of the *hSyn* promoter and allow assessment of their activity via a head mounted mini-microscope (Fig 1A). We prepared 18 mice (9 with injection of GCaMP6 into the RTN and 9 with injection into the RVLM, Fig 1B) and assessed their responses to hypercapnia before and after induction of SE (during chronic phase of epilepsy; 3, 5 and 7-weeks post-SE). After exclusion of 3 RTN-injected mice (Fig 1B) for insufficient fluorescence signals, we analysed the remaining mice for their neuronal responses to hypercapnia before the induction of SE and at 3, 5 and 7-weeks following SE induction with intrahippocampal kainic acid (KA) injection (Fig 1C and Supplementary Fig 1-2). We observed the same range of firing patterns in response to hypercapnia in both the RTN and RVLM as we have previously reported in the RTN (Bhandare *et al*., 2022). These were as follows:

**Figure 2:**
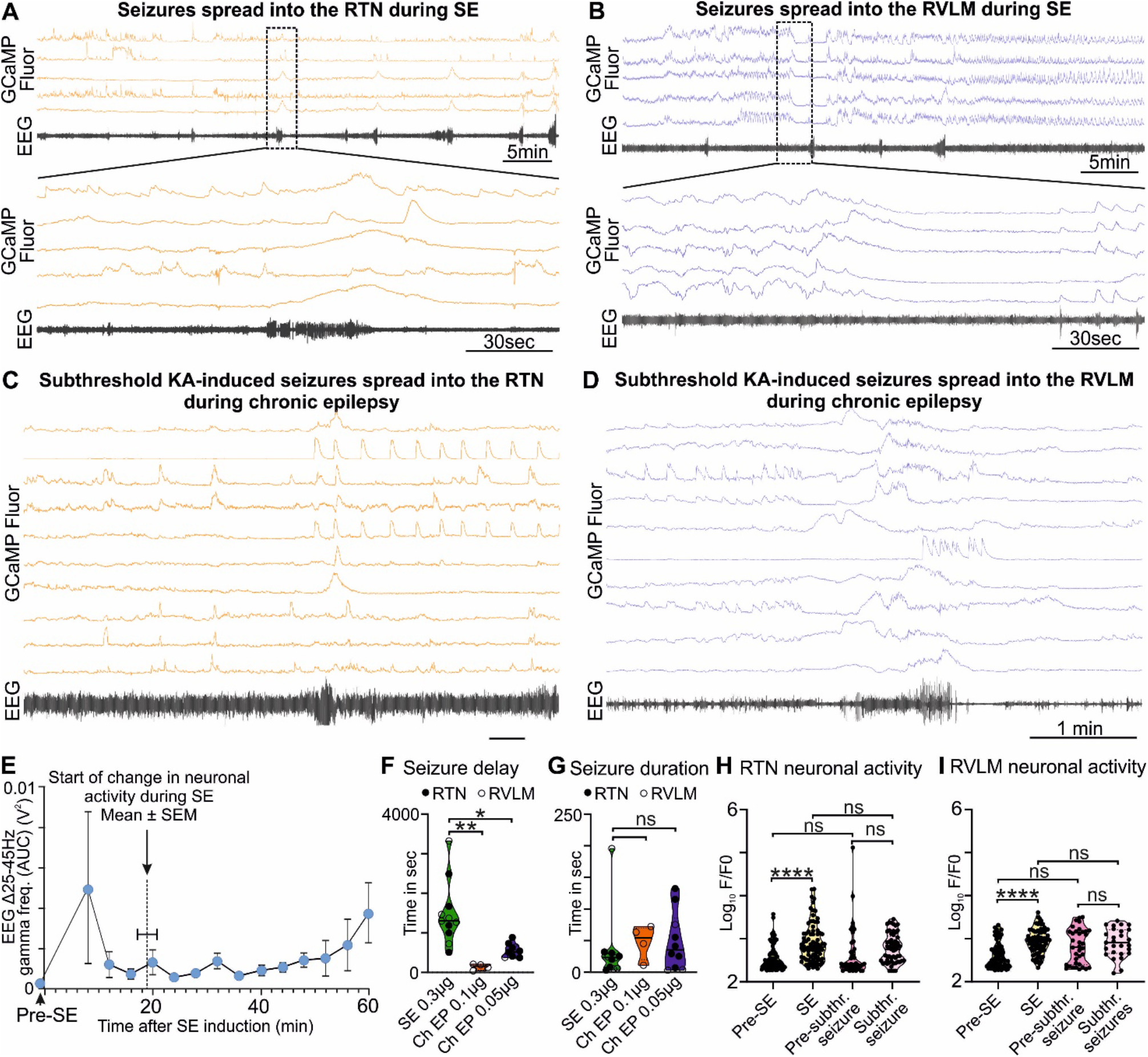
KA-induced seizures alter the activity of brainstem cardiorespiratory neurons. A) KA-induced seizures during status epilepsy (SE) spread into the RTN nuclei. The dotted square shows changes in activity of RTN neurons in freely behaving mice time matched with EEG seizure and expanded under it. Seizure activity after intrahippocampal injection of KA time matched with the activity of RTN neurons (orange). B) KA-induced seizures during status epilepsy spread into the RVLM nuclei. The dotted square shows changes in activity of RVLM neurons in freely behaving mice time matched with EEG seizure and expanded under it. Seizure activity after intrahippocampal injection of KA time matched with the activity of RVLM neurons (blue). The effect of 6- and 3-fold lower dose of intrahippocampal KA (compared with the SE) on C) the RTN and D) the RVLM neuronal activity, respectively, with reduced seizure induction delay time. E) Start of seizure activity i.e., changes in amplitude of gamma-range frequency (25-45Hz) before induction of SE (pre-SE) and after intrahippocampal injection of KA (n=8). Increase in the gamma range frequency (25– 45 Hz) in EEG is characteristic property of KA-induced seizures (Gurbanova *et al*., 2008; Bhandare *et al*., 2016). The seizure start time is compared with the time of first visual aberrant cardiorespiratory neuronal activity calculated from respective Ca^2+^ traces from mice recorded for RTN and RVLM (n=14). The start of change in neuronal activity is represented as a dotted line. During chronic epilepsy in mice seizure delay (F) and duration time (G) with 3- and 6-fold lower dose of intrahippocampal KA (against SE) is compared with times at SE. n=11 at SE (0.3 µg of intrahippocampal KA), and n=4 and 10 (0.1 and 0.05 µg of intrahippocampal KA, respectively) during chronic epilepsy. The average change in activity of all RTN (H) and RVLM (I) neurons during induction of SE and subthreshold KA-induced seizures in comparison with their pre-level (baseline) activities i.e. before induction of SE and before induction of subthreshold KA-induced seizures during chronic epilepsy (at week-7 post-SE). Data in the panel F-I are median and quartile and shown as continuous and dotted line, respectively. P values derived from one-way ANOVA with multiple comparison and corrected with Dunnett’s multiple comparisons test for panel F-G and Brown-Forsythe test for the panel H-I. F* (DFn, DFd) are 2.796 (2, 22) for panel-F, 0.3038 (2, 22) for panel-G, 12.68 (3.000, 150.6) for panel-H, and 39.70 (3.000, 106.7) for panel I. The significant P values are \**p* < 0.05, \*\**p* < 0.01, \*\*\**p* < 0.001 and \*\*\*\**p* < 0.0001. subthr., subthreshold.

*Inhibited (I)*: Displayed spontaneous Ca^2+^ activity at rest, which was greatly reduced during both 3 and 6% inspired CO_2_ and could exhibit rebound activity following the end of the hypercapnic episode.

*Excited - adapting (E_A_)*: Silent or with low level activity at rest and showed the greatest Ca^2+^ activity in response to a change in 3% inspired CO_2_. Following an initial burst of activity, they were either silent or displayed lower-level activity throughout the remainder of the hypercapnic episode. These cells often exhibited rebound activity following the end of the hypercapnic episode.

*Excited - graded (E_G_):* Silent or with low level activity at rest and displayed an increase in Ca^2+^ activity at 3% inspired CO_2_ with a further increase in activity at 6% that returned to baseline upon removal of the stimulus.

*Tonic (T):* Displaying spontaneous Ca^2+^ activity throughout the recording that was unaffected by the hypercapnic episode.

*Non-coding (NC):* Displayed low frequency or sporadic Ca^2+^ activity that had no discernible relationship to the hypercapnic stimulus or any respiratory event.

We were unable to record neuronal activity during spontaneous seizures and all the neuronal responses shown here are from evoked seizures and during response to hypercapnia in mice. This is because the low frequency of spontaneous seizures and their unpredictable occurrence makes it difficult to record the neuronal activity during these seizures. Ca^2+^ imaging cannot be performed continuously due to bleaching of GCaMP6. Thus, it is not possible to systematically record neuronal activity during these rare seizures. Neurons recorded in the RTN were targeted with the GRIN lens (Fig 1D), imaged in freely behaving mice (Fig 1E) and identified by their position relative to the facial (7^th^) nucleus and co-staining for neuromedin B (NMB) (Fig 1F). Neurons in the RVLM were targeted with the GRIN lens (Fig 1G), imaged in freely behaving mice (Fig 1H) and identified by their position relative to the nucleus ambiguus and immunoreactivity for phenylethanolamine N-methyltransferase (PNMT) (Fig 1I).

During SE-induced seizures (recorded via EEG electrodes), large amplitude Ca^2+^ signals could be observed in neurons of both the RTN (Fig 2A, Supplementary Fig 3A, Supplementary Movie 1) and RVLM (Fig 2B, Supplementary Fig 4A, Supplementary Movie 2). This aberrant neuronal activity commenced around 18.9 ± 1.5 minutes after injection of KA (Fig 2E; there was ∼ 6 min delay between intrahippocampal injection and the start of recording) and lasted throughout the period of seizures until this was terminated by ketamine and diazepam injection. We could not record the EEG before injection of intrahippocampal KA as the position of hippocampal cannula was very close to the EEG headmount and this would have necessitated anesthetizing the mice twice (for connection of the EEG electrodes pre-SE and during intrahippocampal KA injection). Due to this technical reason we did not record the EEG prior to intrahippocampal KA injection, however, as shown previously (Cavalheiro *et al*., 1982; Gurbanova *et al*., 2008; Bhandare *et al*., 2016), we found that the seizures had already started at 6 min after KA injection (increase in the power of the gamma-range frequency (25-45Hz) compared to pre-seizure EEG activity (EEG activity during pre-SE hypercapnic response; Fig 2E). During SE induction through intrahippocampal injection of KA (0.3 μg/mouse), we recorded first seizure (EEG burst) with EEG electrodes at 1481.8 ± 240.5 sec (mean ± SEM) after KA injection (Fig 2F) and the seizure lasted for 33.7 ± 16.4 sec (Fig 2G).

### Increased sensitivity and reduced latency to seizure induction with aberrant brainstem neuronal activity in mice with chronic epilepsy

7.9 ± 0.2 weeks after induction of SE (i.e. during the chronic phase of epilepsy), the seizures induced by the lower doses of KA (0.1 μg of KA/mouse, n=4, and 0.05 μg of KA/mouse, n=10) were able to invade the RTN (Fig 2C, Supplementary Movie 4, 0.05 μg of KA) and RVLM (Fig 2D, Supplementary Movie 3, 0.1 μg of KA) and cause aberrant neuronal activity within these nuclei (Supplementary Fig 3B and 4B). We found that the time for the induction of the first seizure/visual EEG burst was significantly reduced compared to the onset of seizures during SE (142.0 ± 37.7 sec for 0.1 μg KA, p = 0.0008, Fig 2F, Supplementary Movie 3 (RVLM imaging) and 552.8 ± 53.5 sec for 0.05 μg KA, p = 0.0016, Fig 2G, Supplementary Movie 4 (RTN imaging)) but there was no change in the duration of the resulting seizures (47.8 ± 13.5 sec for 0.1 μg KA, p = 0.8460, and 48.6 ± 14.5 sec for 0.05 μg KA, p = 0.7190; Fig 2F-G) as compared with SE.

**Figure 3:**
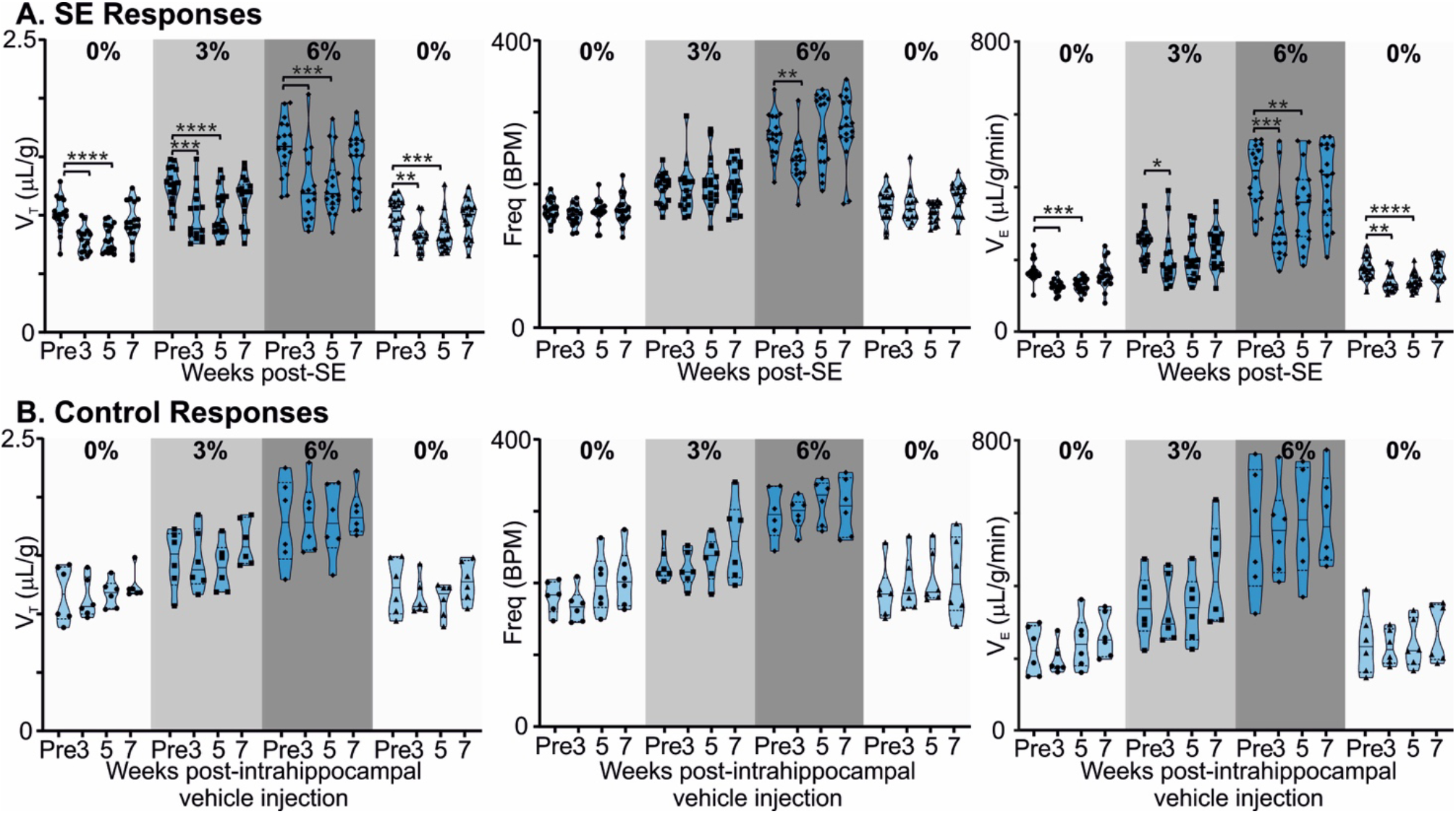
Chronic epilepsy induces a breathing phenotype in mice. A) Changes in tidal (V_T_), frequency (f_R_) and minute ventilation (V_E_) in response to 3 and 6% CO_2_ before and 3, 5 and 7-weeks after induction of SE in freely behaving mice (n=18). Median and quartile are shown with continuous and dotted line, respectively. B) Changes in tidal (V_T_), frequency (f_R_) and minute ventilation (V_E_) in response to 3 and 6% CO_2_ before and 3, 5 and 7-weeks after injection of intrahippocampal PBS in freely behaving mice (control; n=6). Data are median and quartile and shown as continuous and dotted line, respectively. P values derived from two-way repeated measure (mixed effects) ANOVA with Tukey’s multiple comparison are \**p* < 0.05, \*\**p* < 0.001, \*\*\**p* < 0.001 and \*\*\*\**p* < 0.0001.

**Figure 4:**
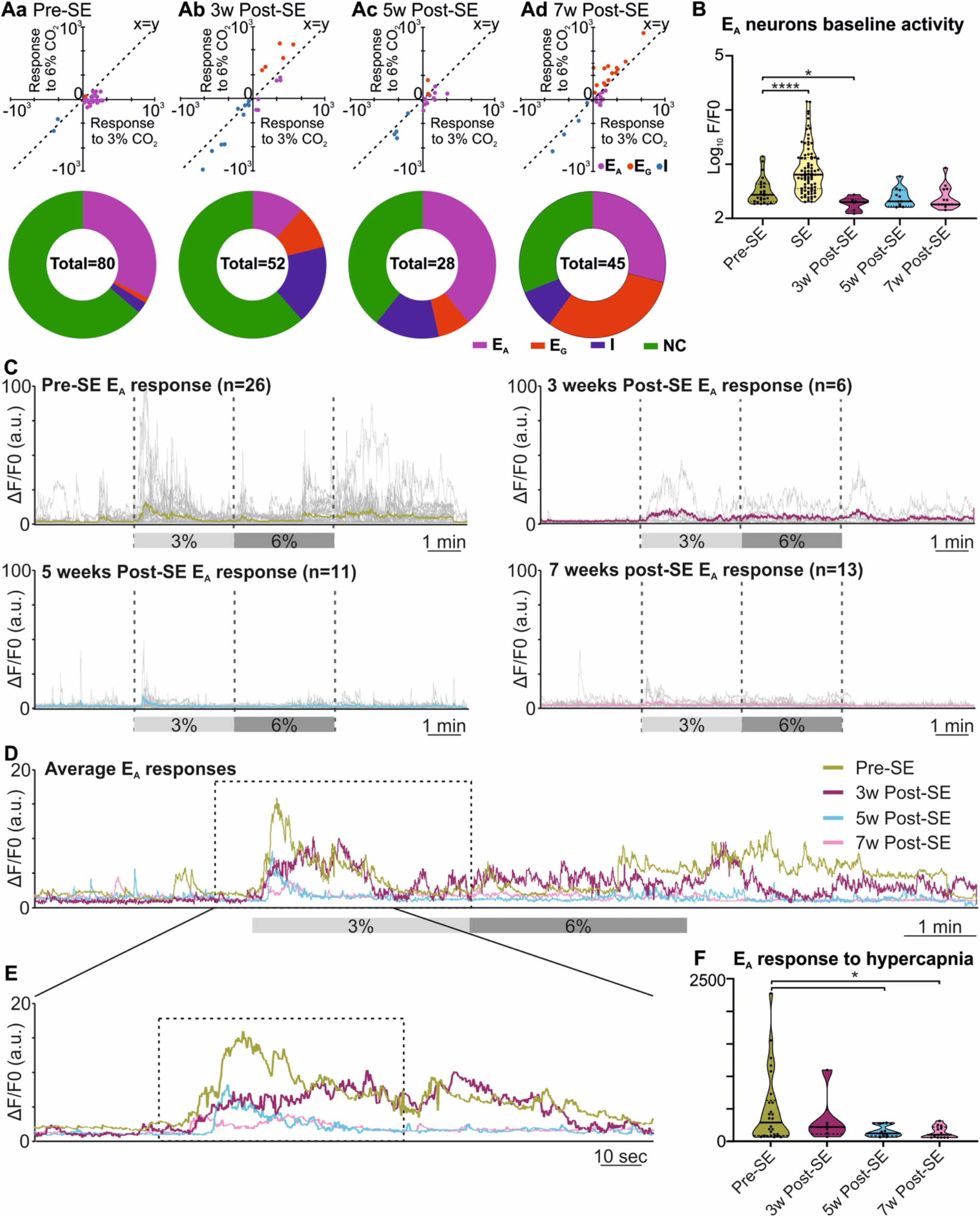
Altered RTN chemosensitive neuronal response to hypercapnia during chronic epilepsy. A) Two component analysis of neuronal categorisation of RTN cells at pre-SE (a) and 3- (b), 5- (c), and 7-weeks (d) post-SE. The change in Ca^2+^ signal (measured as area under the curve) from baseline (room air) elicited by 6% CO_2_ plotted against the Ca^2+^ response elicited by 3% CO_2_. The line of identity (x=y) is shown. E_A_ neurons fall below this line, whereas E_G_ neurons fall above this line and I neurons are predominantly in the negative quadrant of the graph. The donut charts represent the proportion of different types of neuronal responses to hypercapnia in the RTN at pre-SE (a), and 3- (b), 5- (c), and 7-weeks (d) post-SE. B) Baseline activities of RTN Ad neurons at pre-SE, and 3-, 5- and 7-weeks post-SE and compared with neuronal responses during SE. C) RTN neurons individual and average adapting responses to hypercapnia before and 3, 5 and 7-weeks after induction of epilepsy in freely behaving mice. n is number of neurons recorded from 6 mice. In 7-weeks post-SE panel, for technical reasons the recording of recovery from hypercapnia is absent. D) RTN neurons average adapting responses compared between pre and 3, 5 and 7-weeks post-SE. The dotted square shows changes in adapting response of RTN neurons in freely behaving mice and expanded in panel E. E) Changes in RTN neurons adapting response to 3% CO_2_ (dotted box) with pre-hypercapnic baseline activity. F) RTN neurons adapting response to 3% CO_2_ (from the dotted box in panel E) measured as the area under the curve (AUC). Data in the panel B and F are median and quartile and shown as continuous and dotted line, respectively. For comparison of E_A_ vs non-E_A_ neurons in panel-A chi-squared test is used with the false discovery rate procedure for the correction. One-way ANOVA with multiple comparison and Brown-Forsythe correction is used for the comparison in panel B and F. F* (DFn, DFd) are 51.64 (4.000, 110.4) for panel-B and 4.869 (3.000, 21.21) for panel F. P values are \**p* < 0.05 and \*\*\*\**p* < 0.0001.

We compared the baseline activity of all RTN and RVLM neurons before induction SE with activity during SE, and at week 7 post SE (before subthreshold KA induced seizures) with the activity during subthreshold KA induced seizures in chronic epilepsy (Fig 2H-I). The activity of all neurons in both nuclei was significantly increased at SE compared to pre-SE (p <0.0001 for both RTN and RVLM) but did not change at subthreshold KA-induced seizures compared to their pre-levels (p = 0.4512 and 0.9163 for the RTN and RVLM, respectively). Most importantly, between RTN and RVLM groups, there was no difference in the activity levels at SE and lower dose of KA-induced seizures compared to pre-SE and subthreshold KA-induced seizures, respectively (Fig 2H-I). The baseline activity in both nuclei, when recorded for all neurons, showed no difference between the pre-SE level and the level recorded at 7-weeks post SE (p = 0.9716 and 0.0662 for the RTN and RVLM, respectively; Fig 2H-I).

Our findings show that the activity of the cardiorespiratory brainstem neuronal network, although far from the site of seizure origin, can still be altered by the subthreshold (3 and 6-fold lower) dose of KA-induced seizures in the hippocampus during the chronic phase of epilepsy. This plausibly suggests a mechanism for the cardiorespiratory failure during SUDEP: aberrant neuronal activity in the RTN and RVLM during and immediately after a seizure. The equal magnitude of brainstem neuronal responses to SE and subthreshold dose of KA during chronic epilepsy (p = 0.3152 and 0.9992 for the RTN and RVLM, respectively) suggests an increased sensitivity and susceptibility of brainstem cardiorespiratory neuronal network to seizures during chronic epilepsy that could plausibly arise from continued invasion of spontaneous seizures into the brainstem nuclei (Fig 2H-I).

### Development of a compromised breathing phenotype in mice with chronic epilepsy

The RTN is an important nucleus that contributes to the chemosensory control of breathing. By contrast the RVLM projects to the sympathetic and parasympathetic preganglionic neurons and controls cardiovascular parameters such as heart rate and blood pressure. RVLM neurons also exhibit chemosensitivity (Koganezawa & Paton, 2014). As seizures invade these nuclei, we expected that there might be altered chemosensory control during the development of chronic epileptic states following induction of SE. We therefore examined how responses to hypercapnia in mice subjected to intrahippocampal KA injection or saline injection (control, surgical intervention but no SE induction) changed with time (Fig 3). We evaluated hypercapnic neuronal and breathing responses from week-3 post-SE in mice that were already experiencing spontaneous seizures and compared them with pre-SE responses. As the mice were already experiencing spontaneous seizure at week-3 post-SE, this means our post-SE neuronal and breathing responses were outside classical latent period (Levesque & Avoli, 2013; Lévesque *et al*., 2016; Rusina *et al*., 2021).

In mice with chronic epilepsy, the baseline tidal volume (V_T_) during breathing of room air was significantly lowered when assessed 3 and 5-weeks after induction of SE (p <0.0001; Fig 3A). The increases in V_T_ observed during mild (3% CO_2_) and moderate (6% CO_2_) hypercapnia were also depressed compared to their values pre-SE (p = 0.0004 and <0.0001 for 3% CO_2_ at 3-and 5-weeks post-SE, respectively, and p = 0.0003 and 0.0001 for 6% CO_2_ at 3- and 5-weeks post-SE, respectively; Fig 3A). At week 7, epileptic mice showed partial recovery of reduced V_T_ responses to hypercapnia (p = 0.3644 and 0.3155 at 3% and 6% CO_2_, respectively, compared to pre-SE levels; Fig 3A). Interestingly the respiratory frequency was not altered by induction of SE or during the establishment of chronic epilepsy. As might be expected, the changes in minute ventilation (V_E_) reflected the changes apparent in V_T_ (Fig 3A).

By contrast, in control mice, the ventilatory responses to hypercapnia were maintained in control mice over the entire study period (Fig 3B). V_T_ in baseline (room air breathing) became stronger at 7 weeks post intrahippocampal saline injection (p = 0.83; Fig 3B). The lung inspiratory capacity of mice increases with age with maximum change happening between 3-12 months (Huang *et al*., 2007; Schulte *et al*., 2019). An increasing baseline breathing trend observed in the control mice is attributable to this maturation (Fig 3B). The age of control and epileptic mice at the pre-SE/PBS recording was between 90-170 days and at 7 weeks post-SE/PBS recording it was between 145-230 days. There was no significant age difference between two groups at pre (p = 0.66) and 7-weeks post-SE (p = 0.21) recordings (unpaired t test with Welch’s correction). Following induction of SE the normal maturational increase in baseline breathing was not observed (compare Fig 3A and B), which could be due to effect of SE or development of chronic epilepsy in these mice. Overall, our data shows that the alteration in baseline breathing and the sensitivity of breathing to different levels of CO_2_ evident in the epileptic mice are due to the induction of epilepsy rather than the surgical intervention.

### Attenuated responses to hypercapnia during chronic epilepsy by adapting RTN chemosensitive neurons

Given that the adaptive ventilatory changes to hypercapnia altered following induction of SE, we tested whether there might also be changes in the activity of chemosensory neurons in the RTN. Previously we have documented that RTN neurons exhibit a range of firing patterns during hypercapnia (Bhandare *et al*., 2022) and in the present study we also observed similar patterns of neuronal responses to hypercapnia from RTN neurons both before and after induction of SE (Fig 1B and Supplementary Fig 1). Our categorisation of RTN neurons into the E_A_, E_G_, I, and NC subtypes was supported by a two-component analysis in which we plotted the change in Ca^2+^ activity (from baseline) elicited by 6% inspired CO_2_ versus the change in Ca^2+^ activity elicited by 3% inspired CO_2 (_Fig 4 Aa-d x-y plots). As before (Bhandare *et al*., 2022), we found that most frequent neuronal response to hypercapnia before the induction of SE was an excited adapting (E_A_) pattern with an increase in activity immediately following the increase in inspired CO_2_ (Figs 1B and 4Aa donut chart).

Following induction of SE, changes in the pattern of RTN neuronal responses to hypercapnia became apparent over the following weeks (Fig 4). We performed the chi-squared test and corrected with the false discovery rate procedure for multiple comparisons (Curran-Everett, 2000) and found that the proportion of E_A_ neurons (vs non-adapting subtypes) was significantly decreased at week 3 (week 3 vs pre-SE, χ^2^ = 7.54, *p* = 0.006; week 5 vs pre-SE, χ^2^ = 0.42, *p* = 0.51; week 7 vs pre-SE, χ^2^ = 0.17, *p* = 0.68; Fig 1B and 4Aa-d donut charts). We noticed the similar trend when we looked at the baseline Ca^2+^ activity of E_A_ neurons (Fig 4B). Although the area under curve (AUC) of the F/F_0_ in the baseline was significantly increased during SE compared to pre-SE (Fig 4B, p<0.0001), at 3 weeks after SE this baseline activity was decreased compared to pre-SE level (Fig 4B, *p* = 0.03194). At later time points the baseline activity recovered to pre-SE levels (Fig 4B). By contrast to the E_A_ neurons, we saw an increase in the proportion of E_G_ and I neurons after induction of SE in these mice with greatest increase in number of E_G_ neurons being evident at week 7 (Fig 4Aa-d donut charts). This suggests considerable plasticity in the chemosensory networks of the RTN.

Because E_A_ neurons were the major subtype in our recordings, we further quantified the adapting response by averaging the individual changes in GCaMP6 fluorescence at each monitoring point (Fig 4C-F). This showed that the adapting Ca^2+^ signal was weaker at 3, 5 and 7-weeks following induction of SE compared to before SE (Fig 4D-E). As a further quantification, we measured the area under the curve (AUC) of the adapting response during 3% inspired CO_2_ (Fig 4F). This again confirmed the diminished neuronal activation during hypercapnia at week 5 and 7 post-SE (*p* = 0.016 and 0.011, respectively) (492.4 ± 109.0 at pre-SE compared to 337.4 ± 155.3, 155.6 ± 25.9 and 139.0± 24.9 at 3, 5, and 7-weeks post-SE, respectively). Overall, there was a transient reduction in the baseline E_A_ neuronal activity in the RTN at three weeks, and also a sustained reduction in the amplitude of the responses of the E_A_ neurons to hypercapnia that was still evident 7 weeks post-SE induction.

### Responses of RVLM neurons to hypercapnia during chronic epilepsy

Neurons of the RVLM exhibited similar types of responses to hypercapnia as those in the RTN (Fig 1B) and this was confirmed by a two-component analysis of the neuronal responses (Fig 5Aa-d x-y-plots). As for the RTN neurons, before induction of SE the adapting response was most common (Fig 5Aa donut chart). Post-SE RVLM neuronal responses to hypercapnia showed a difference in the proportion of E_A_ neurons compared to non-adapting that was only evident at week 7 when tested with the chi-squared test and corrected with the false discovery rate procedure for multiple comparison (week 3 vs pre-SE χ^2^ = 0.3, *p* = 0.58; week 5 vs pre-SE, χ^2^ = 2.58, *p* = 0.1; week 7 vs pre-SE χ^2^ = 7.2, *p* = 0.007). There was also an increased proportion of inhibited and graded neurons at week 5 and 7 (Fig 5Aa-d donut charts). Baseline Ca^2+^ activity of RVLM adapting neurons (AUC of the F/F_0_) did not change during development of chronic epilepsy, but as expected it showed a significant increase during induction SE (p <0.0001; Fig 5B). Averaging the GCaMP6 fluorescence activity of individual adapting RVLM neurons showed that these responses remained constant even after induction of SE (Fig 5C-F). However, the magnitudes of all RVLM responses (F/F_0_) were less than those of the RTN neurons (compare Figs 4C-F with Figs 5C-F). The AUC analysis of RVLM E_A_ neurons confirmed that their responses to hypercapnia remained stable at all timepoints after induction of SE (Fig 5F, AUC for RVLM adapting responses were 197.0 ± 24.0 at pre-SE compared to 123.2 ± 18.5, 163.2 ± 48.6 and 138.2 ± 24.8 at 3, 5 and 7-weeks post-SE, respectively).

**Figure 5:**
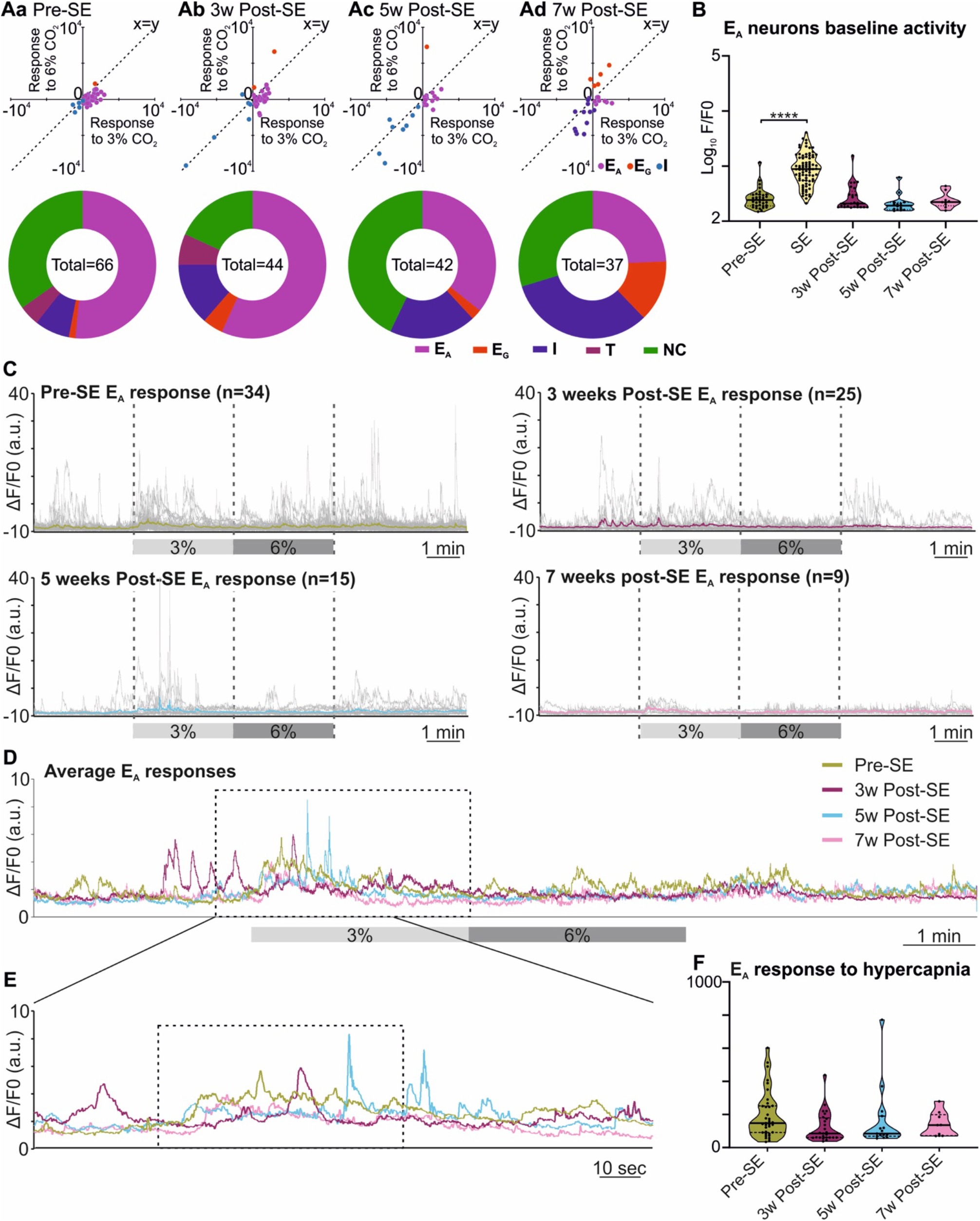
Reponses of neurons in the RVLM are maintained following induction of SE. A) Two component analysis of neuronal categorisation of RVLM cells at pre-SE (a) and 3- (b), 5- (c), and 7-weeks (d) post-SE. The change in Ca^2+^ signal (measured as area under the curve) from baseline (room air) elicited by 6% CO_2_ plotted against the Ca^2+^ response elicited by 3% CO_2_. The line of identity (x=y) is shown. E_A_ neurons fall below this line, whereas E_G_ neurons fall above this line and I neurons are predominantly in the negative quadrant of the graph. The donut charts represent the proportion of different types of neuronal responses to hypercapnia in the RVLM at pre-SE (a), and 3- (b), 5- (c), and 7-weeks (d) post-SE. B) Baseline activities of RVLM E_A_ neurons at pre-SE, and 3-, 5- and 7-weeks post-SE and compared with neuronal responses during SE. C) RVLM neurons individual and average adapting responses to hypercapnia before and 3, 5 and 7-weeks after induction of epilepsy in freely behaving mice. n is number of neurons recorded from 9 mice. D) RVLM neurons average adapting responses compared between pre and 3, 5 and 7-weeks post-SE. The dotted square shows changes in adapting response of RVLM neurons in freely behaving mice and expanded in panel E. E) RVLM neurons adapting response to 3% CO_2_ (dotted box) with pre-hypercapnic baseline activity. F) RVLM neurons adapting response to 3% CO_2_ (from the dotted box in panel E) measured as the AUC. Data in the panel B and F are median and quartile and shown as continuous and dotted line, respectively. For comparison of E_A_ vs non-E_A_ neurons in panel-A chi-squared test is used with the false discovery rate procedure for the correction. One-way ANOVA with multiple comparison and Brown-Forsythe correction is used for the comparison in panel B and F. F* (DFn, DFd) are 81.34 (4.000, 113.7) for panel-B and 1.651 (3.000, 38.93) for panel F. P value is \*\*\*\**p* < 0.0001.

### Induction of SE and development of spontaneous seizures in RTN and RVLM group

Chemoconvulsant-induced models of chronic epilepsy are variable with respect to development of SE and subsequent development of spontaneous seizures (Lévesque *et al*., 2016). Hence, to evaluate these parameters in both RTN and RVLM group of mice, we performed series of quantitative analyses. Firstly, we quantified the Racine behavioural scale values for each mouse and compared these between RTN and RVLM group with chi-squared test and corrected with the false discovery rate procedure for multiple comparison, which showed no difference between two groups (Fig 6A; scale-1 χ^2^ = 2.7, *p* = 0.09; scale-2 χ^2^ = 0.9, *p* = 0.33; scale-3 χ^2^ = 0.25, *p* = 0.61; scale-3.5 χ^2^ = 2.0, *p* = 0.16; scale-4 χ^2^ = 1.39, *p* = 0.23; scale-4.5 χ^2^ = 1.1, *p* = 0.29; scale-5 χ^2^ = 4.2, *p* = 0.04 (but this was not significant as it was above the threshold *p* value corrected for multiple comparisons by the false discovery rate procedure - 0.0062); scale-6 χ^2^ = 0.76, *p* = 0.38).

**Figure 6:**
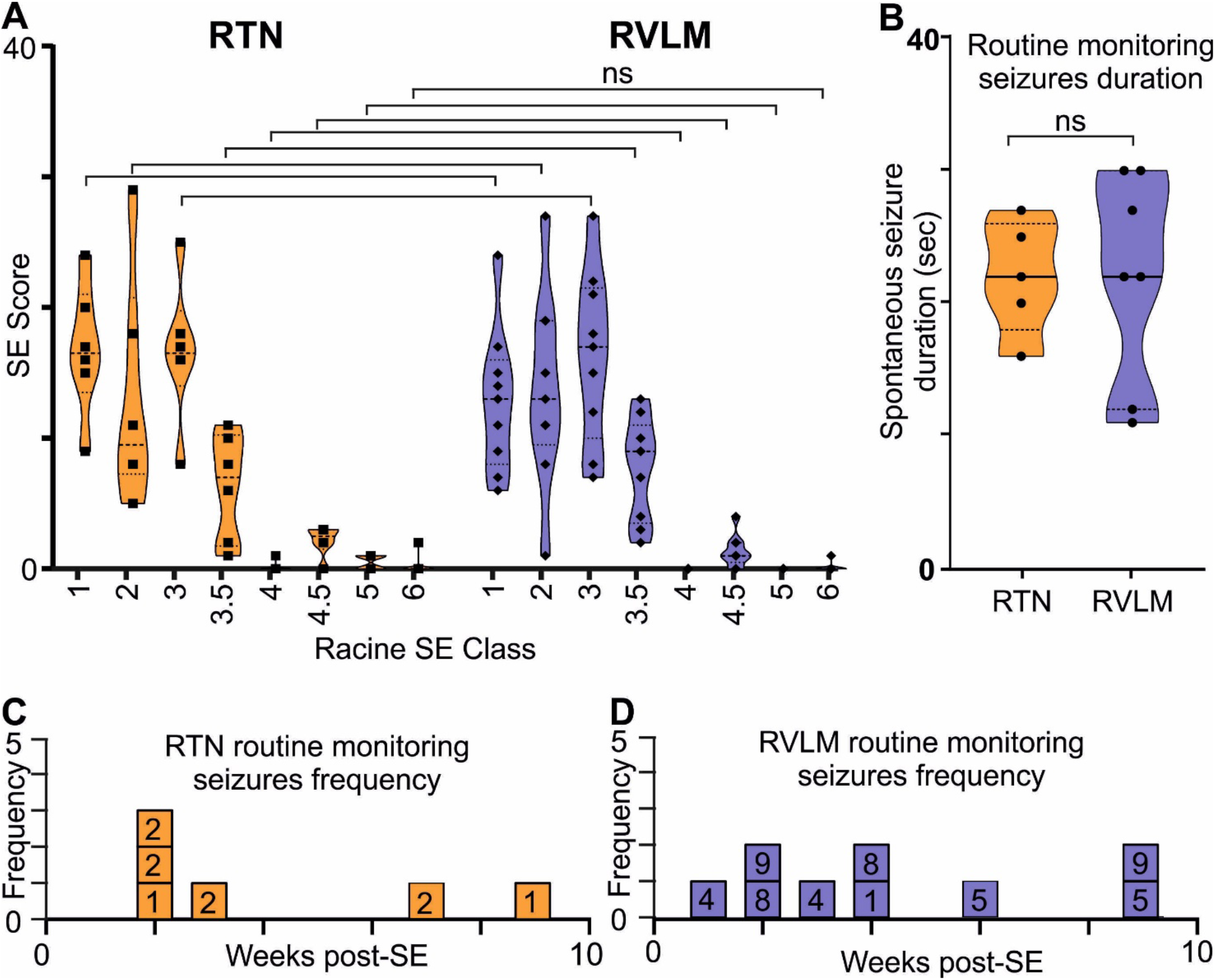
A comparable induction of SE and development of spontaneous seizures in RTN and RVLM group of mice. A) Comparison of Racine behavioural scale score during induction of SE in mice recorded for RTN and RVLM. x-axis, Racine behavioural class and y-axis, number (score) of specific SE behaviour. B) Comparison of duration of spontaneous seizures noted during routine monitoring and manually recorded in individual mice from RTN and RVLM group of mice. Weekly spontaneous seizures frequency noted (during routine mouse monitoring), and video recorded in individual mice recorded for C) RTN and D) RVLM neuronal activity after induction of SE. The numbers in the boxes in C and D are mouse identifiers and correspond to the identifying numbers in the supplementary data figures. Data in panel A and B are median and quartile and shown as continuous and dotted line, respectively. For comparison of Racine score between RTN and RVLM chi-squared test is used with the false discovery rate procedure for the correction. Mann-Whitney nonparametric t-test is used for the comparison in panel B.

Secondly, the duration and frequency of spontaneous seizures noted during routine animal monitoring was not different between RTN and RVLM (Fig 6B-D). To test whether the routine monitoring (30-45min every day) was insufficient to capture the seizure behaviour of the mice, we performed longer term continuous video monitoring in three additional RTN mice (6.9hrs/day/mouse for 15 days over 7 weeks, Fig 7A-B). All three mice experienced spontaneous seizures with similar timing (between weeks 2 and 4 after SE) to the cohort of mice reported in Fig 6.

**Figure 7:**
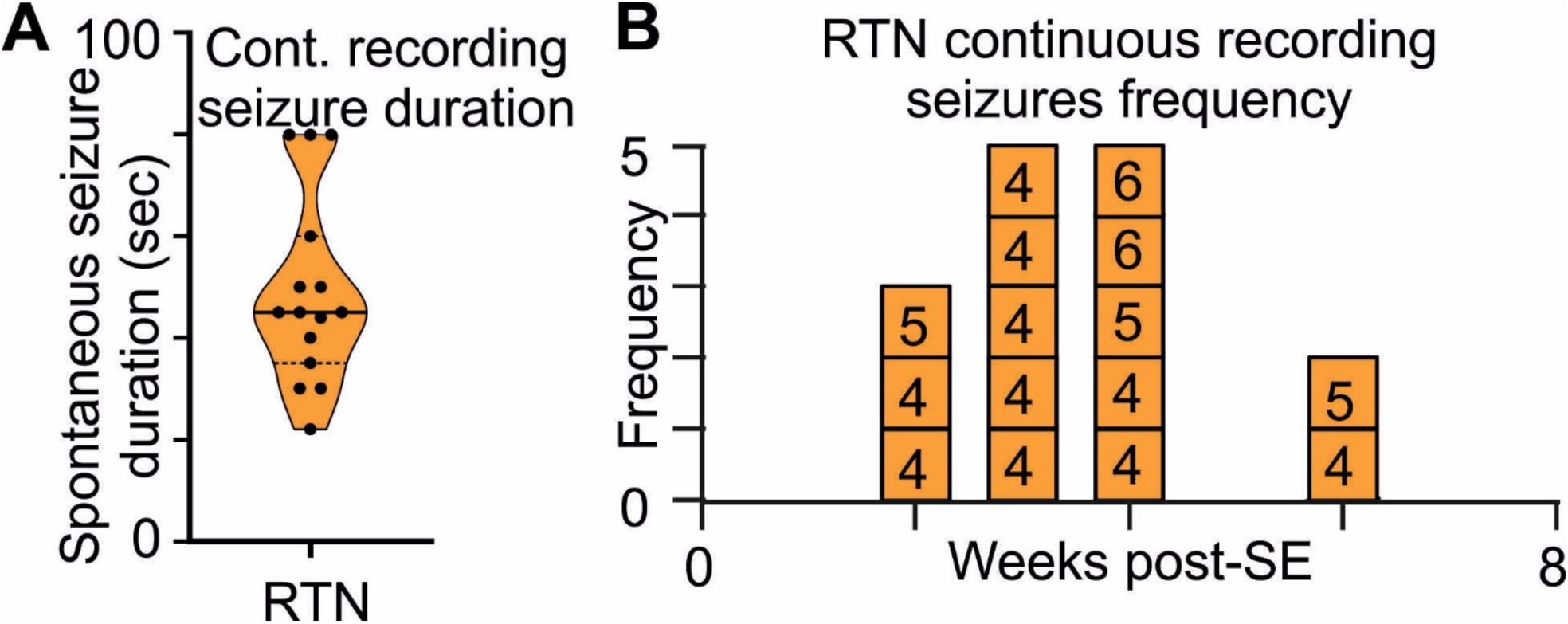
Long-term monitoring of spontaneous seizures in RTN mice. A) Duration and B) frequency of spontaneous seizures recorded in longer-term video monitoring of 3 mice selected for recordings from RTN neurons. Data in panel A is median and quartile and shown as continuous and dotted line, respectively. The numbers in the boxes in B are mouse identifiers and correspond to the identifying numbers in the supplementary data figures.

The routine monitoring was performed during the daily check-up of the mice over the 7 weeks post SE and amassed approximately 30hrs/mouse of visual inspection. This is some 3.5 times shorter than the longer-term video recording presented in Fig 7. However as both methods gave results with similar numbers of seizures normalized to total recording time, we conclude that the routine daily monitoring was sufficient to sample the number, intensity and duration of seizures exhibited by the mice.

Thirdly, the seizure delay time (time to induce first electrographic seizure after KA injection) was decreased during chronic epilepsy in both RTN and RVLM mice (compared to SE seizures delay time) when we injected 6-fold lower dose of KA (dose-dependently). Electrographic seizures are seizures that are evident on EEG with or without associated noticeable behavioural signs. Importantly, seizure duration and delay time was not different between both RTN and RVLM group of mice during both SE and chronic seizures (Fig 2F-G).

Finally, analysis of neuronal Ca^2+^ activity during induction of SE showed that the average activity of the RTN and RVLM neurons was increased to similar level (p<0.0001; Figs 4B and 5B) suggesting that both nuclei had similar effect of induction of SE. Additionally, we compared all neuronal activities after injection of KA during induction of SE and subthreshold KA-induced seizures and these were also not different between RTN and RVLM mice (Fig 2H-I). This suggests that both the RTN and RVLM mice had an equal SE trigger and development of comparable spontaneous seizures. Therefore, the lack of effect of spontaneous seizures on the responses of RVLM adapting neurons to hypercapnia is unlikely to be because the SE or spontaneous seizures were weaker in these mice.

## Discussion

Modelling of SUDEP is experimentally challenging due to its unpredictable nature. While we have not directly modelled SUDEP, we microinjected a chemoconvulsant into a restricted brain region to induce localised acute and chronic spontaneous seizures. This approach allowed us to discern the effects of seizures on the central cardiorespiratory system that could contribute to SUDEP. In our study, chronic spontaneous seizures are defined as behavioural seizures as we did not record concurrent EEG activity in these mice. The chemoconvulsant induced rodent models of epilepsy show a classical latent period after injection of chemoconvulsant, which is associated with reconfiguration of neuronal network, neuronal loss, reactive gliosis, inflammation and neurogenesis before the start of spontaneous seizures (Leite *et al*., 2002; Buckmaster, 2004). The latent period varies from model to model but in intrahippocampal KA injection in mice, the latent period lasts on average between 4 and 30 days after KA administration (Levesque & Avoli, 2013; Lévesque *et al*., 2016; Rusina *et al*., 2021). In our study, mice were experiencing spontaneous seizures from week 2 after injection of KA. Therefore, the neuronal and breathing changes that are reported at 3, 5 and 7 post-SE are outside the classical latent period and is the effect of SE and spontaneous seizures. More importantly, as mice were in the plethysmograph chamber for 30-60 min under observation before any recordings commenced, this excluded the possibility that the observed changes in neuronal activity and breathing were the effect of an immediately preceding seizure. However, we cannot exclude the possibility of seizures occurring a few hours or days before recording as we did not perform 24 hrs video recording to confirm this.

The number of neurons recorded from session to session that we recorded from the same mouse varied. This is because movement artefacts (which may differ from session to session) sometimes prevented clear resolution of activity of some neurons. We also encountered these issues in our previous study where we imaged neurons in the same region (Bhandare *et al*., 2022). The reasons for this are probably that the brainstem is extracranial and more mobile (hence the recordings are more susceptible to movement artefacts).

Adaptive increases in tidal volume (V_T_) or breathing frequency (f_R_) in response to hypercapnia (chemosensory reflex) are an important component of normal central respiratory homeostasis (Spyer & Gourine, 2009; Guyenet, 2014). Mice with chronic epilepsy developed a breathing phenotype with reduced V_T_ which was evident at 3 and 5-weeks post-SE. In chronic epileptic mice, baseline breathing in room air was reduced compared to the naïve state and their chemosensory adaptive responses to hypercapnia were significantly decreased at mild (3%) and moderate (6%) CO_2_. This reduced respiratory breathing phenotype in mice at week 3 and 5 post-SE has its neuronal correlate in the RTN adapting neuronal responses. Our findings are in line with a previous study where 30 days after induction of SE, rats showed a reduced breathing response (Maia *et al*., 2020). Similarly, people with epilepsy showed lower hypercapnic ventilatory responses correlated with higher postictal rise in CO_2_ following generalized convulsive seizures (Sainju *et al*., 2019). Interestingly, at week-7 post-SE we noticed a partial recovery of breathing phenotype in mice with chronic epilepsy possibly because of the emergence of neurons in the RTN that exhibited graded responses to hypercapnia. This suggests considerable adaptive plasticity in the respiratory neuronal networks as has been shown in other contexts (Ling *et al*., 2001; Dahan *et al*., 2007).

By imaging neuronal Ca^2+^ activity at cellular resolution in key respiratory and cardiovascular brainstem nuclei in awake mice before, during and after the establishment of the chronic epileptic state, we have avoided one of the major drawbacks of prior studies: the use of anaesthesia which unavoidably alters circuit and cellular brain function. Clinical and preclinical findings have suggested that forebrain seizures might spread to the medullary neuronal network and cause central cardiorespiratory arrest (Aiba & Noebels, 2015; Dlouhy *et al*., 2015). Our real-time cellular imaging of activity of individual neurons in the RTN and RVLM does indeed demonstrate that seizures spread into both nuclei and cause aberrant neuronal activity in these critical networks during and following the period of the seizure. This further strengthens the findings by Aiba and Noebels (Aiba & Noebels, 2015) by establishing the effect of seizures on specific subsets of brainstem cardiorespiratory autonomic neurons. We have shown recently that the RTN neuronal responses do not change in mice due to their age or any other factor and in fact remain same across weeks of interimaging interval (Figure 1-figure supplement 6 of ref (). Therefore, the change in the activity of brainstem neurons that we see after induction of SE is the effect of seizures rather than an intrinsic time dependent process in the brainstem.

Our findings extend the observation of reduced hypercapnic ventilatory response in both animal models and people with epilepsy (Sainju *et al*., 2019) by providing further insight into: 1) a potential underlying mechanism indicated by the alterations of neuronal responses in the RTN, a major chemosensory nucleus; and 2) suggest that the central respiratory networks have sufficient plasticity to at least partially compensate for compromised breathing responses in epilepsy.

In addition to the ability of seizures to invade acutely into the RTN and RVLM nuclei, we found that the chronic epileptic state caused long-lasting alterations in the ability of RTN neurons to respond to hypercapnia. Adapting responses to hypercapnia evident in the naïve state (before induction of SE) were greatly attenuated when tested 3, 5 and 7 weeks following establishment of the epileptic state (Fig 4C-F). This correlated with reduced hypercapnic and normal room air breathing responses at week 3 and 5 post-SE in these mice. On the other hand, proportion of excitatory graded RTN neurons increased at week-7 post SE, which correlates with evidence of partial recovery of hypercapnic breathing response at 7 weeks in these mice.

SUDEP is defined as sudden and unexpected death of person with epilepsy, who is otherwise healthy. One third of people with epilepsy do not respond to currently available anti-epileptic medications and are more vulnerable to SUDEP. Furthermore, the incidence of SUDEP is higher in people with drug-resistant epilepsy (Tomson *et al*., 2008; Ryvlin *et al*., 2013). Case studies have found that the poor seizure control, particularly of generalised tonic-clonic seizures (Walczak *et al*., 2001), a history of any seizure in last three months before SUDEP (compared to none) (Langan *et al*., 2005) and increased frequency of tonic-clonic seizures per year (Nilsson *et al*., 1999) are significant risk factors for SUDEP.

Despite the correlative link between seizure history and SUDEP, its mechanism remains unclear. Although findings from human SUDEP cases and animal studies establish a central role for cardiac arrhythmia (Bateman *et al*., 2010; Bhandare *et al*., 2017), altered inter-ictal heart rate variability (Surges *et al*., 2009), cardiac arrest (Ryvlin *et al*., 2013), ventricular fibrillation (Naggar *et al*., 2014), bradycardia (Kalume *et al*., 2013), terminal apnoea (Ryvlin *et al*., 2013), obstructive respiratory apnoea (Jefferys *et al*., 2019; Irizarry *et al*., 2020), respiratory arrest (Faingold *et al*., 2010), ictal/postictal hypoventilation (Bateman *et al*., 2010) as the proximal cause of SUDEP, the specific neuronal circuits involved, and the reasons for why their dysfunction leads to cessation of breathing or cardiovascular activity are unknown. This makes prevention and treatment of SUDEP more difficult. Our longitudinal imaging of neurons pre- and post-SE induction enabled us to document the effects of repeated chronic spontaneous seizures, a major risk factor for SUDEP, on the function of cardiorespiratory neurons and hypercapnic breathing responses. The weakening of the chemosensory reflex via loss of chemosensitivity in the RTN could plausibly increase the risk of SUDEP. For example, we speculate that if breathing were to cease due to invasion of spreading depolarisation and silencing of electrical activity into the brainstem, arterial blood would become hypercapnic and acidic. Ordinarily, this would act as a powerful stimulant to restart breathing once the spreading depolarisation had passed. However, the weakened chemosensory responses of RTN neurons that we observed in the chronic epileptic state might make it harder to restart breathing and could thus increase the risk of SUDEP. This speculation needs to be tested in a more specific animal model of SUDEP, e.g. DBA/1 mice where the incidence of respiratory arrest following audiogenic seizures increases with seizure repetition (Faingold *et al*., 2010) and DBA/2 where mice have a 100% incidence of respiratory arrest depending on their age. Although these mice die from obstructive apnoea, this likely arises from reduced muscle tone and respiratory related central drive that serves to dilate the airways during inspiration (Irizarry *et al*., 2020).

While we have not studied the effect of seizures on chemosensory neurons of the raphe nucleus, agonists for serotonin receptors (a key raphe neurotransmitter) have a protective effect in blocking seizure-induced SUDEP in DBA mice (Faingold *et al*., 2011; Ma *et al*., 2022) and mice with genetic deletion of serotonin neurons (Buchanan *et al*., 2014). Raphe neurons project to the RTN (Wu *et al*., 2019) and the pre-Bötzinger complex (Ma *et al*., 2022) and contribute to chemosensory responses in these nuclei. Thus, activation of serotonin receptors could help to compensate for the weakened responses of the chemosensitive RTN neurons and enhance the stimulatory effect of CO_2_ on breathing (Mulkey *et al*., 2007; Wu *et al*., 2019). This could be also the likely mechanism of partial recovery of breathing responses at week-7 post SE and emergence of graded response in the RTN. Nevertheless, the action of serotonin could also be mediated through its direct effect on seizure activity (Buchanan *et al*., 2014; Schoonjans *et al*., 2017), arousal (Buchanan & Richerson, 2010), sleep-wake regulation and circadian rhythm (Miyamoto *et al*., 2012) and warrants further investigation.

In contrast to altered RTN neuronal responses to hypercapnia during chronic epilepsy, RVLM neurons showed no change in their hypercapnic responses after SE. Findings from epilepsy monitoring units (So *et al*., 2000; Bateman *et al*., 2010; Ryvlin *et al*., 2013; Mueller *et al*., 2014; Dlouhy *et al*., 2015) and preclinical studies (Faingold *et al*., 2010; Jefferys *et al*., 2019; Maia *et al*., 2020) indicate that respiratory arrest rather than cardiac failure is the major contributor to SUDEP. Our findings are thus consistent with this, as they show a greater immediate effect of repeated seizures on the RTN chemosensory neurons than on the RVLM neurons. Although, either central respiratory or obstructive apnoea contributes to SUDEP, it sets in train a sequence of events that include hypoxemia and acidosis that could indirectly trigger failure of the central cardiovascular system and eventual cardiac arrest (Ryvlin *et al*., 2013).

A final implication of our findings is that longitudinal monitoring of respiratory chemosensitivity, combined with other established clinical risk factors, might be a useful biomarker test to identify people at risk of SUDEP. Although a pre-epileptic baseline for respiratory chemosensitivity could not be established in people with epilepsy, regular measurement of chemosensitivity via a simple non-invasive breathing test, which can be readily performed in epilepsy monitoring units (Sainju *et al*., 2019), could establish trends in chemo-responsiveness to indicate increased or diminished risk of SUDEP.

## Materials and Methods

Experiments were performed in accordance with the European Commission Directive 2010/63/EU (European Convention for the Protection of Vertebrate Animals used for Experimental and Other Scientific Purposes) and the United Kingdom Home Office (Scientific Procedures) Act (1986) with project approval from the University of Warwick’s AWERB (PP1674884).

### Viral handling

We used AAV-9-syn-GCaMP6s vector with synapsin promoter (pGP-AAV-syn-GCaMP6s-WPRE.4.641 at a titre of 1x10^13^ GC·ml^-1^, Addgene, Watertown, MA, USA), and therefore it transduced neurons showing higher tropism for the AAV 2/9 subtype, and did not transduce non-neuronal cells. The AAV uses the synapsin promoter. Virus was aliquoted and stored at −80°C. On the day of injection, it was removed and stored at 4°C, loaded into graduated glass pipettes (Drummond Scientific Company, Broomall, PA, USA), and placed into an electrode holder for pressure injection.

### Viral transfection of RTN and RVLM neurons

Adult male C57BL/6J mice (8-10 weeks old and 20-30 g) were anaesthetized with isofluorane (4%; Piramal Healthcare Ltd, Mumbai, India) in pure oxygen (4 L·min^-1^). Adequate anaesthesia was maintained with 0.5-2% Isofluorane in pure oxygen (1 L·min^-1^) throughout the surgery. Mice received a presurgical subcutaneous injection of atropine (120 µg·kg^-1^; Westward Pharmaceutical Co., Eatontown, NJ, USA) and meloxicam (2 mg·kg^-1^; Norbrook Inc., Lenexa, KS, USA). Mice were placed in a prone position into a digital stereotaxic apparatus (Kopf Instruments, Tujunga, CA, USA) on a heating pad (TCAT 2-LV: Physitemp, Clifton, NJ, USA) and body temperature was maintained at a minimum of 33°C via a thermocouple. The head was levelled, at bregma and 2 mm caudal to bregma, and graduated glass pipettes containing the virus were placed stereotaxically into either the RTN or RVLM (Fig 1D,G). The RTN was defined as the area ventral to the caudal half of the facial nucleus, bound medially and laterally by the edges of the facial nucleus (coordinates with a 9⁰ injection arm angle: -1.0 mm lateral and -5.6 mm caudal from Bregma, and -5.2 to -5.5 mm ventral from the surface of the cerebellum; Fig 1D). The RVLM defined as a triangular area ventral to the nucleus ambiguus (NA), lateral to the inferior olive (IO) or pyramids (Py) and medial to the spinal trigeminal sensory nucleus (Sp5) (coordinates with a 0⁰ injection arm angle: 1.3 mm lateral and -5.7 mm caudal from Bregma, and -5.15 mm ventral from the surface of the cerebellum; Fig 1G). The virus solution was pressure injected (<300 nL) unilaterally. Pipettes were left in place for 3-5 minutes to prevent back flow of the virus solution up the pipette track. Postoperatively, mice received intraperitoneal (IP) injections of buprenorphine (100 µg·kg^-1^; Reckitt Benckiser, Slough, UK). Mice were allowed 2 weeks for recovery and viral expression, with food and water *ad libitum*.

### Implantation of EEG electrodes, hippocampal cannula and GRIN lens

Mice expressing GCaMP6 were anaesthetized with isofluorane, given pre-surgical drugs, placed into a stereotax, and the head was levelled as described above. Four holes were drilled for implantation of 2 EEG recording, a ground and a reference electrode (Fig 1A). EEG electrodes were PFA-coated silver wire of diameter: 0.254mm (Bilaney Consultants Ltd, UK) (coordinates with a 0⁰ injection arm angle: Recording electrodes- ±1.5 mm lateral and 1.0 mm rostral from Bregma, ground and reference electrodes- ±2.5 mm lateral and - 1.5 mm caudal from Bregma). Silver wires were implanted and secured in position with SuperBond™ (Prestige Dental, Bradford, UK). EEG wires were passed through the pedestal (Bilaney Consultants Ltd, UK), and pedestal was implanted and secured in place over the head with SuperBond™. Another two holes were drilled each for implantation of unilateral hippocampal cannula (Fig 1A and C, coordinates with a 0⁰ injection arm angle: 1.8 mm lateral and -2.0 mm caudal from Bregma, and -1.6 mm ventral from the surface of the dura) and GRIN lens (coordinates as above). Stainless steel cannula, 26 gauge and 10mm long, was implanted in the hippocampus and secured in place with SuperBond™. To widen the lens path whilst producing the least amount of deformation of tissue, a graduated approach was taken; firstly a glass pipette was inserted down the GRIN lens path to a depth 500 µm above where the lens would terminated and left in place for 3 mins; this procedure was then repeated with a blunted hypodermic needle. The GRIN lens (600 µm diameter, 7.3 mm length; Inscopix, Palo Alto, CA, USA) was then slowly inserted at a rate of 100µm·min^-1^ to a depth ∼1300 µm above the target site, then lowered at a rate of 50µm·min^-1^ to a depth ∼300 µm above the RTN or RVLM target region (coordinates as above with 300 µm above the target site). The lens was then secured in place with SuperBond™. Postoperatively, mice received buprenorphine, and were allowed 2 weeks for recovery, with food and water *ad libitum*.

### Baseplate installation

Mice expressing GCaMP6 and implanted with EEG electrodes, hippocampal cannula and GRIN lens were anaesthetized with isofluorane, given pre-surgical drugs, and placed into a stereotax as described above. To hold the miniaturized microscope during recordings, a baseplate was positioned over the lens and adjusted until the cells under the GRIN lens were in focus. The baseplate was then secured with superbond™, and coated in black dental cement (Vertex Dental, Soesterberg, the Netherlands) to stop interference of the recording from ambient light. Mice were allowed 1 week for recovery, with food and water *ad libitum*.

*Induction of status epilepsy (SE) with EEG recording and Ca^2+^ imaging in freely moving mice* All mice were trained with dummy camera and habituated to plethysmography and flexi glass open field chamber before imaging. Instrumented mice were anaesthetized with isofluorane and placed into a stereotax as described above. Mice EEG electrodes were connected to wires and miniature microscope with integrated 475 nm LED (Inscopix, Palo Alto, CA, USA) was secured into the baseplate. SE was induced in mice with total of 0.3 μg (in 50nl volume) of unilateral kainic acid (KA) injection through hippocampal cannula with Hamilton Neuros syringe (Model 75 RN, Essex Scientific Laboratory Supplies Ltd., UK). Mice were taken off the anaesthesia and placed into open field chamber. The EEG recording and Ca^2+^ imaging were started ∼6 min after KA injection. Mice were scored every minute for their behavioural seizures using Racine scale (1- Rigid posture or immobility; 2- Stiffened, extended, and often arched (Straub) tail; 3- Partial body clonus, including forelimb or hindlimb clonus or head bobbing; 3.5- Whole body continuous clonic seizures while retaining posture; 4- Rearing; 4.5- Severe whole body continuous clonic seizures while retaining posture; 5- Rearing and falling; 6- Tonic–clonic seizures with loss of posture or jumping). At 60 minutes or recording 1-2 of class 5/6 scale seizure following injection of KA (whichever happened earlier), mice were anaesthetized with isofluorane and SE was terminated with intraperitoneal injection of ketamine (50 mg/kg) and diazepam (20 mg/kg).

During SE, EEG activity was recorded, amplified and filtered using the NeuroLog system (Digitimer, Welwyn Garden City, UK) connected to a 1401 interface and acquired on a computer using *Spike2* software (Cambridge Electronic Design, Cambridge, UK). The EEG activity raw data were DC removed. The power in the “gamma” frequency range (25– 45 Hz) was analysed, as previously shown (Gurbanova *et al*., 2008; Bhandare *et al*., 2016). A power spectrum analysis of EEG (AUC, V^2^) was done from 2 min blocks taken every 4 min apart after the start of recording. Video data was recorded with *Spike2* software and was synchronised with the EEG activity (Supplementary Movie 1-2). GCaMP6 fluorescence was visualised during SE through the GRIN lens, using nVista 2 HD acquisition software (Inscopix, Palo Alto, CA, USA). Calcium fluorescence was optimised for each experiment so that the histogram range was ∼150-600, with average recording parameters set at 10 frames/sec with the LED power set to 10-20 mW of light and a digital gain of 1.0-4.0. A TTL pulse was used to synchronize the calcium signalling to the behavioural data and EEG trace. All images were processed using Inscopix data processing software (Inscopix, Palo Alto, CA, USA). GCaMP6 movies were ran through preprocessing algorithm (with spatial and temporal downsampling), crop, spatial filter algorithm (0.005 - 0.5 Hz), motion correction and cell identification through CNMF-E (Constrained Nonnegative Matrix Factorization for microEndoscopic data) analysis to generate the identified cell sets. Cell sets were imported into *Spike2* software for processing. All Ca^2+^ traces from specific time point were averaged and analysed for area under the curve (AUC) between the start 3% hypercapnia and 180 sec after it.

### Plethysmography

Mice were placed into a custom-made 0.5 L plethysmography chamber, with an airflow rate of 1 l·min^-1^. The plethysmography chamber was heated to 31^0^C (thermoneutral for C57/BL6 mice). CO_2_ concentrations were sampled via a Hitech Intruments (Luton, UK) GIR250 Dual Sensor Gas analyzer or ML206 gas analyzer (ADinstruments, Sydney, Australia) connected to the inflow immediately before entering the chamber. The analyser had a delay of ∼15-20 sec to read-out the digital output of gas mixture. Pressure transducer signals and CO_2_ measurements were amplified and filtered using the NeuroLog system (Digitimer, Welwyn Garden City, UK) connected to a 1401 interface and acquired on a computer using *Spike2* software (Cambridge Electronic Design, Cambridge, UK). Video data was recorded with *Spike2* software and was synchronised with the breathing trace. Airflow measurements were used to calculate: tidal volume (V_T_: signal trough at the end of expiration subtracted from the peak signal during inspiration, converted to mL following calibration), and respiratory frequency (*f*R: breaths per minute). Minute ventilation (V_E_) was calculated as V_T_ x *f*R.

### Hypercapnia in freely behaving mice

Mice were tested for hypercapnic challenge before and 3, 5 and 7-weeks after induction of SE. Instrumented mice were allowed ∼30 mins to acclimate to the plethysmograph. The LED was activated through a TTL pulse synchronised with the *Spike2* recording and 3 min of baseline recordings were taken (gas mixture: 0% CO_2_ 21% O_2_ 79% N_2_). The mice were then exposed to 3 min epochs of hypercapnic gas mixture at different concentrations of CO_2_ (3, and 6% in 21% O_2_ balanced N_2_). Following exposure to the hypercapnic gas mixtures, CO_2_ levels were reduced back to 0% and calcium signals were recorded for a further 4 minutes recovery period.

### Seizure threshold in mice with chronic epilepsy

After recording 3, 5 and 7-weeks post-SE hypercapnia responses and during chronic phase of epilepsy (8-weeks post-SE), mice were tested for seizure threshold. Instrumented mice were anaesthetized with isofluorane, placed into a stereotax and connected to EEG electrodes and miniature microscope as described above. Mice were injected through intrahippocampal cannula with 3 or 6-fold lower dose of KA (0.1μg, n=4 and 0.05μg, n=10 of KA/mouse) compared to SE. Mice were taken off the anaesthesia, placed into open field chamber and EEG recording and Ca^2+^ imaging were started as stated above. Mice were recorded until first EEG seizure burst was recorded (Supplementary Movie 3-4) and seizures were terminated as above.

### Statistical analysis

Statistical analysis was performed in GraphPad Prism software (version 9.1.0). Statistical significance for changes in breathing was determined using two-way repeated measure ANOVA followed by t tests with the Tukey’s correction and shown as a violin plot. Statistical analysis of the changes in RTN and RVLM neuronal adapting hypercapnic response, baseline activity and changes during SE and subthreshold KA-induced seizures were performed via the Brown-Forsythe and Welch ANOVA with Dunnett T3 correction. This alternative test was used as the data were not normally distributed and the variance differed between groups in the comparison, thus invalidating use of a conventional ANOVA. The seizure delay and duration time were analysed using a one-way ANOVA followed with the Dunnett’s correction. Results are shown as a violin plot with superimposed data points. The Racine seizure scores between two groups (violin plot with superimposed data points) and number of adapting vs non-adapting neuronal responses (donut charts) were compared with chi-squared test and corrected with the false discovery rate procedure for multiple comparisons (Curran-Everett, 2000). Comparisons were done between pre- and post-treatment (SE or control groups). P ≤ 0.05 was considered significant. \*\*\*\**p* ≤ 0.0001, \*\*\**p* ≤ 0.001, \*\**p* ≤ 0.01 and \**p* ≤ 0.05. Results are presented as mean ± SEM throughout the text.

### Immunohistochemistry

Mice were humanely killed by pentobarbital overdose (>100 mg·kg^−1^) and transcardially perfused with paraformaldehyde solution (4% PFA; Sigma-Aldrich, St Louis, MO, USA). The head was removed and postfixed in PFA (4°C) for 3 days to preserve the lens tract. The brains were removed and postfixed in PFA (4°C) overnight. Brainstems or hippocampus were serially sectioned at 50-70 μm. Free-floating sections were incubated for 1 hour in a blocking solution (PBS containing 0.1% Triton X-100 and 5% BSA). Primary antibodies (rabbit anti-Neuromedin-B [NMB; 1:1000; SAB1301059; Sigma-Aldrich, St Louis, MO, USA], or rabbit anti-phenylethanolamine N-Methyltransferase [PNMT; 1:500; 204179-T36-SIB; Stratech, Ely, UK] antibody), or (chicken anti-GFAP [GFAP; 1:500; ab4674; Abcam PLC, Cambridge, UK], or (mouse anti-NeuN [NeuN; 1:500; ab104224; Abcam PLC, Cambridge, UK] were added and tissue was incubated overnight at room temperature.

Slices were washed in PBS (6 × 5 mins) and then the secondary antibodies were added: either donkey anti-rabbit Alexa Fluor 680 (1:250; Jackson Laboratory, Bar Harbor, ME, USA), or donkey anti-rabbit Alexa Fluor 594 (1:250; Jackson Laboratory, Bar Harbor, ME, USA), or donkey anti-mouse Alexa Fluor 680 (1:250; Jackson Laboratory, Bar Harbor, ME, USA) antibody, or donkey anti-chicken Alexa Fluor 594 (1:250; Jackson Laboratory, Bar Harbor, ME, USA) antibody tissue was then incubated for 2-4 hours at room temperature. Tissue was washed in PBS (6 × 5 min). Slices were mounted, coverslipped and examined using a Zeiss 880 confocal microscope with ZEN aquisition software (Zeiss, Oberkochen, Germany).

## Acknowledgements

Funding: This work was supported by the Epilepsy Research UK (ERUK) Project Grant and Wellcome-Warwick QBP funds. ND was a Royal Society Wolfson Research Merit Award Holder and AB is an ERUK Emerging Leader Fellow. We thank Prof Mark Wall and Dr Robert Dallmann for their valuable feedback on the manuscript.

## Author contributions

AB designed the study, performed experiments, recordings, immunohistochemistry analysed data and contributed to writing of the manuscript; ND conceived and designed the study, analysed data and contributed to writing of the manuscript.

## Data and Materials availability

All data is presented in the paper. Raw unprocessed data available on request.

---

**Supplementary Figure 1:**
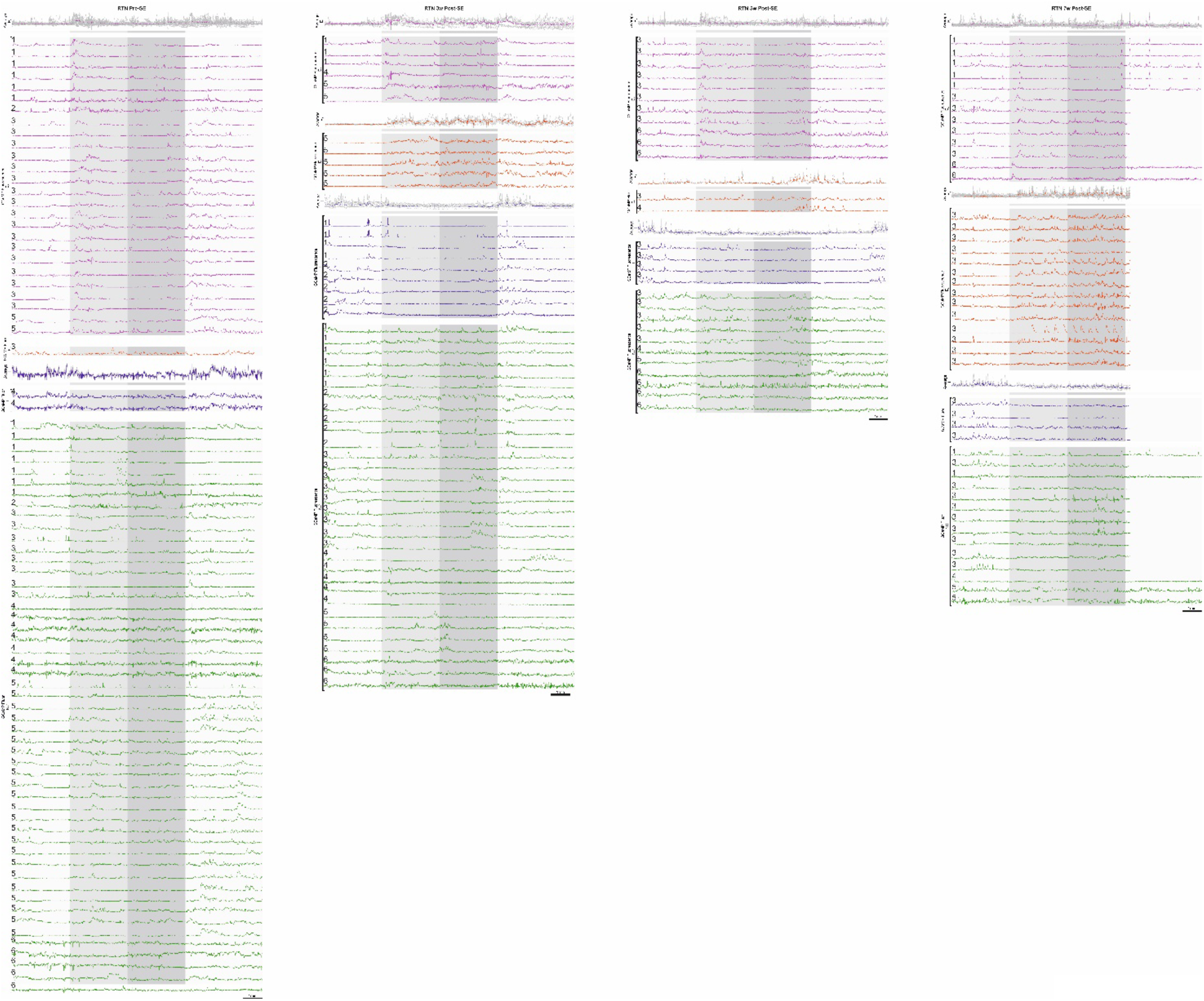
RTN neurons’ responses to the hypercapnic challenge at different time points in epilepsy. RTN excited graded (E_G_), excited adapting (E_A_), inhibited (I), and non-coding (NC) neuronal responses, at pre-SE, and 3-, 5- and 7-weeks post-SE, time-matched with hypercapnia (light grey-3% CO_2_, medium grey-6% CO_2_) and average waveforms of E_G_, E_A_, and I. Animal number is indicated on the left hand side of the neuronal trace. In week-7 post-SE RTN neurons, for technical reasons the recording of recovery from hypercapnia is absent.

**Supplementary Figure 2:**
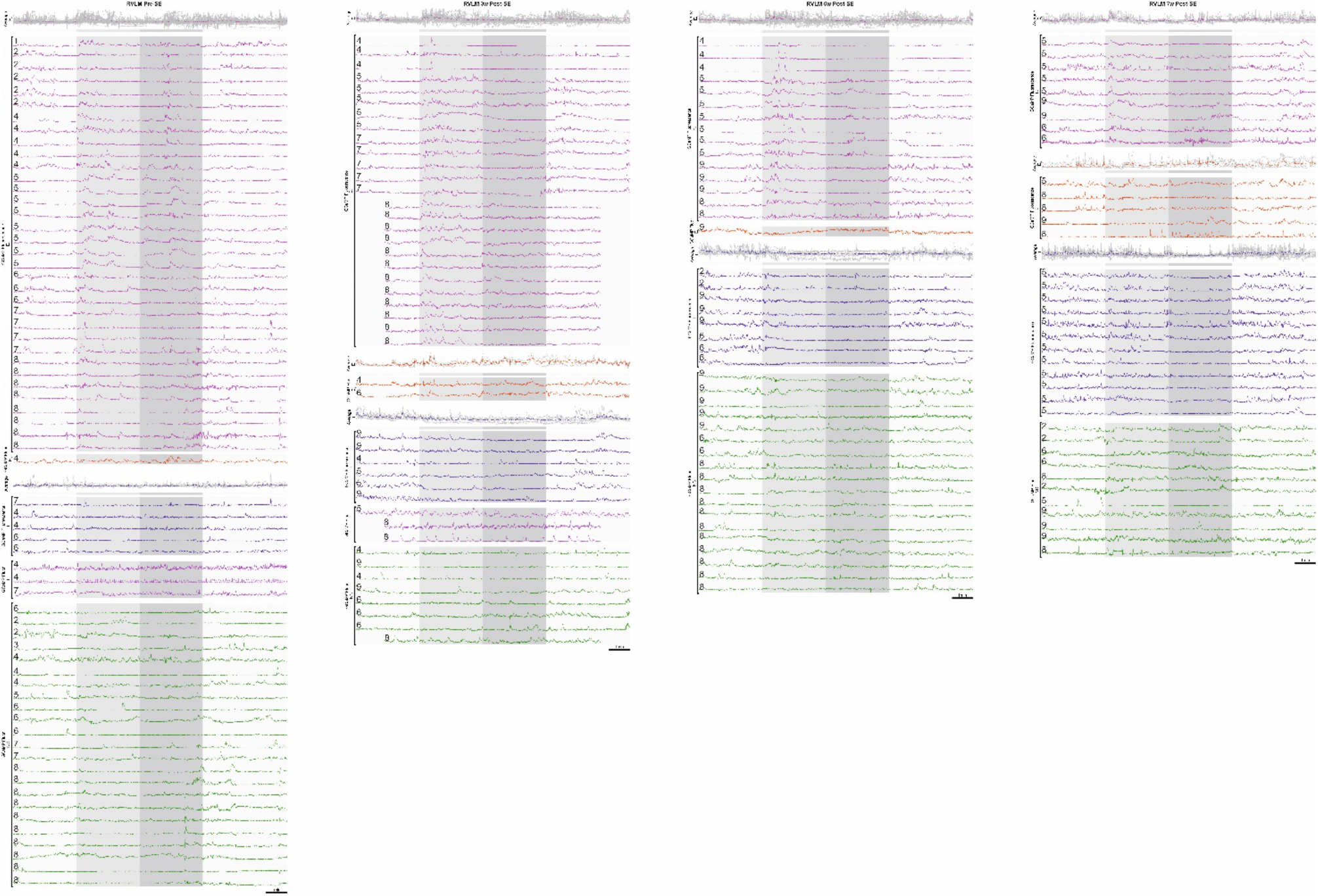
RVLM neurons’ responses to the hypercapnic challenge at different time points in epilepsy. RVLM excited graded (E_G_), excited adapting (E_A_), inhibited (I), tonic (T), and non-coding (NC) neuronal responses, at pre-SE, and 3-, 5- and 7-weeks post-SE, time-matched with hypercapnia (light grey-3% CO_2_, medium grey-6% CO_2_) and average waveforms of E_G_, E_A_, and I. Animal number is indicated on the left hand side of the neuronal trace.

**Supplementary Figure 3:**
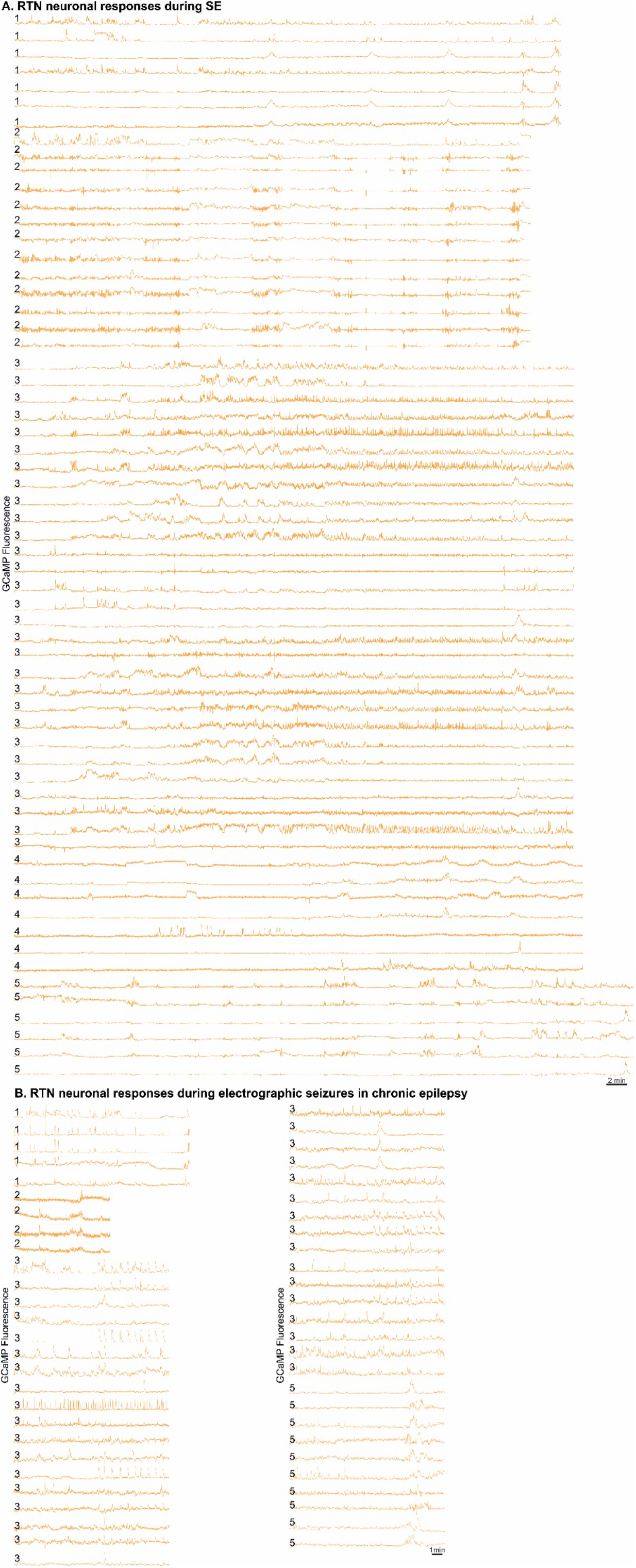
RTN neurons’ activity during KA induced SE and subthreshold KA-induced seizures during chronic epilepsy. Animal number is indicated on the left hand side of the neuronal trace.

**Supplementary Figure 4:**
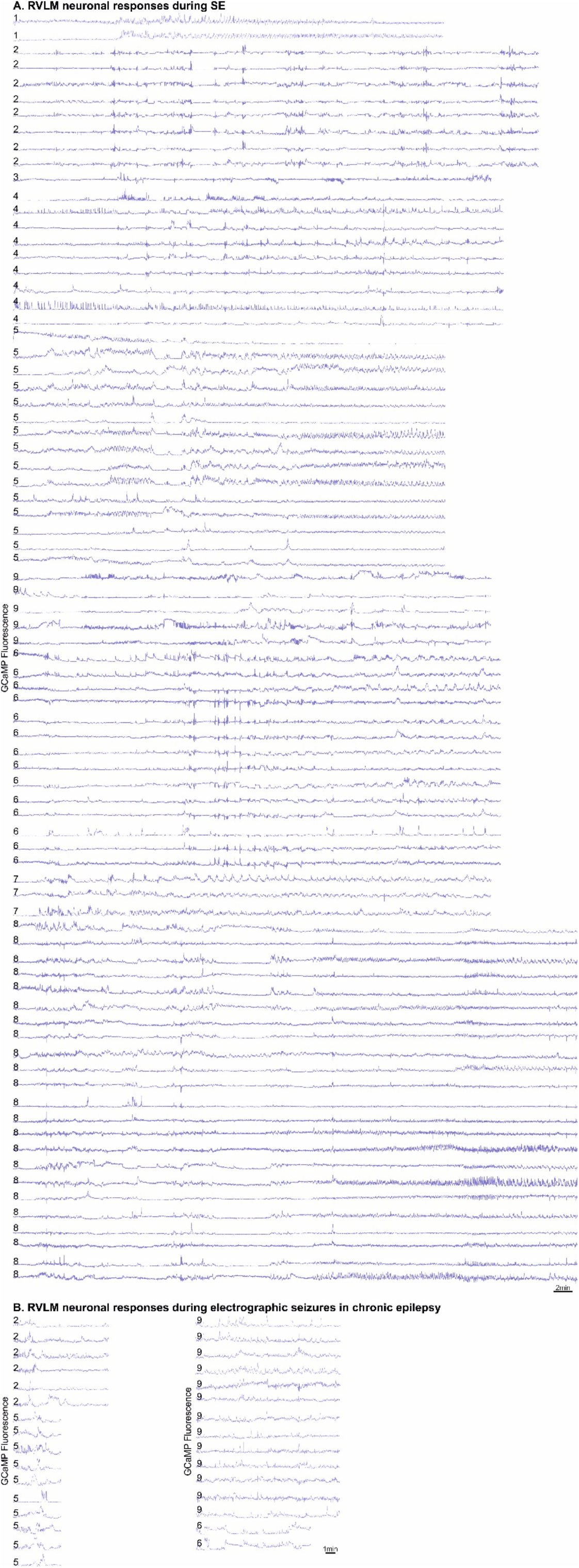
RVLM neurons’ activity during KA induced SE and subthreshold KA-induced seizures during chronic epilepsy. Animal number is mentioned on the left hand side of the neuronal trace.

**Supplementary Movie 1:** Seizures spread into the RTN neurons and disturb their activity during induction of SE via intrahippocampal KA injection in freely behaving mice. Movie is 8x fast forwarded. Selective neuronal traces are shown and matched with the ROIs drawn around GCaMP6s fluorescent neuronal cell bodies.

**Supplementary Movie 2:** Seizures spread into the RVLM neurons and disturb their activity during induction of SE via intrahippocampal KA injection in freely behaving mice. Movie is 8x fast forwarded. Selective neuronal traces are shown and matched with the ROIs drawn around GCaMP6s fluorescent neuronal cell bodies.

**Supplementary Movie 3:** During chronic epilepsy, 3 times of lower dose of KA (0.1 μg) compared to SE showed reduced latency for induction and spread of seizures into the RVLM neurons. Movie is 8x fast forwarded. Selective neuronal traces are shown and matched with the ROIs drawn around GCaMP6s fluorescent neuronal cell bodies.

**Supplementary Movie 4:** During chronic epilepsy, 6 times of lower dose of KA (0.05 μg) compared to SE showed reduced latency for induction and spread of seizures into the RTN neurons. Movie is 8x fast forwarded. Selective neuronal traces are shown and matched with the ROIs drawn around GCaMP6s fluorescent neuronal cell bodies.

